# Differences in personality / temperament in the endangered tortoise *Testudo hermanni* and their implications on dispersal and conservation

**DOI:** 10.1101/2021.08.05.455297

**Authors:** Alexandra Rodriguez, Sébastien Caron, Jean Marie Ballouard

## Abstract

Behavioural studies are more an more implicated in species conservation. Determining individuals personality in the case of reintroduction operations may be very useful. Actually, indiviuals temperament may be associated to their dispersal capacities and their habilities to adapt to novel environments. Considered as asociable species, few studies have been conducted on reptiles and this is even worse in the case of endangered species. Hermann tortoise, Testudo hermanni, an endemic species from Mediterranean region is endangered because of the lost and modification of its habitats. Before conducting reintroduction actions it is important to have more information on individuals personality traits. We have tested the reaction of three groups of tortoises (domestic, wild and from the SOPTOM center) when confronted to a novel environment and to human presence. The aim was to discriminate individuals with « bold » and « shy » behaviours. Behavioural profiles are different between the three groups of tortoises, the domestic group appeared to be bolder than the wild one. Moreover, for the wild group, bold individuals travel longer distances in the field, sometimes outside the protected areas. Thus, it is important to take into account the personality of individuals choosen for translocation projects.

## Introduction

Translocations are methods more and more used in wildlife management. They consist in moving animals from one location to another and they are more and more implemented in conservation programs during the last decades. They are increasingly applied as a form of mitigation for endangered species (Germano et al. 2015; Sullivan et al. 2015) and they need to be carefully prepared to optimize the choose of individuals that have more chance to survive in the new places where they are released.

In this context taking into account factors as physiology and behavior becomes more and more important when selecting the individuals and this is particularly true in the case of reptiles’ personality (Germano et al. 2017).

The terms personality and temperament indicate the interindividual differences in the expression of the behavior present early in invididuals’ lives and stable across the time and contexts (Jones 1977, Bates 1989, Goldsmith et al. 1987, Sih et al. 2004).

Temperament and personality can be explored by the study of different behavioral characteristics as sociability, activity, reactivity and emotivity in social, predatory and novel situations (Bates 1986). These behavioral characteristics may condition dispersal behavior, approach towards novel objects and novel food items consumption (Sinn et al. 2008). Thereby, in non-human animals those aspects of behavior may constraint individuals’ travelled distances and home ranges characteristics (Digemanse et al. 2002, Van Oers et al. 2004a). According to several studies conducted in humans, the differences observed in temperament would have genetic and physiologic bases (Emde et al. 1992, Plomin et al. 1993, Groothuis and Carere 2005) and would be modulated by the individuals’ experience, which generates personality. When suites of correlated bahaviours are expressed across the contexts, we talk about syndromes or coping styles (Wechsler 1995, Sih et al. 2004).

In personality literature individuals have been classified as bold or shy depending on the degree of exploratory behaviour’s they express.

Few studies have been conducted on reptiles, however, differences in temperament have been observed in lizards for example and these differences appeared as being involved in learning processes (Carazo et al. 2014). Actually, in this case, bold individuals appeared to learn faster than shy ones because of their tendency to explore earlier the objects and the experimental apparatus. In the Namibian rock agama, *Agama planiceps*, it has been demonstrated that bold individuals are more easily trapped than shy ones (Carter et al. 2012)

Hermann tortoise, *Testudo hermanni*, is an endangered tortoise only present in the Mediterranean region and its geographic distribution has been declining during the last decades as a consequence of modifications in landscape use and structure (Livoreil 2009). Studies on animal personality have been conducted on common, domestic and invasive species (Jones 1977a, Dingemanse et al 2005, Rodriguez 2010, Martin et Fitzgerald 2005) but not on endangered ones. More information is necessary to understand habitat use by those species and exploration of personality traits as exploration of novel environments or antipredatory reactions may be useful in conservation strategies.

The Village des Tortues Center offers the possibility to study *Testudo hermanni* behavior and to compare three different groups with different past experiences. There are almost wild individuals (manipulated very few times), very wild ones, and ‘domestic’ individuals very often manipulated by technicians and with a frequent human as Village visitors come to see them.

We tested novel environment explorative behavior and antipredatory reactions, and we hypothesized that reactions in the tests would differ depending on the individuals’ past experience. We wondered if there would be a correlation between the novel environment behaviors and the antipredatory reactions.

We conducted tests in order to compare personality between males and females and between ‘wild’ and ‘domestic’ individuals. We also tested for the temporal repeatability of the behavioral responses when individuals were confronted two or four times with the novel environment. We classified individuals as bold or shy and characterized these profiles. Finally, for the wild group we measured the distances traveled by the individuals in the nature in order to know if there was a difference in dispersal behavior between bold and shy individuals.

## Methods

### Preliminary experiments

We observed four Hermann tortoises in a novel environment test during 1h in order to define the duration of the study experiments and to identify all the behavioral items that can be observed during this kind of test. We video recorded the preliminary test and all the subsequent experiments with a Canon Power Shot SD A2500 camera. We classified the behaviors in four categories: walk, immobility, visual exploration, olfactive exploration (Table 1). We paid particular attention to visual and olfactive exploration as it has been demonstrated that visual attention in important in novel situations in *Testudo Hermanni* (Chrzanowska et al. 2015).

**Table 1.** Observed behaviors.

**Observed behaviors**
| **Behaviors** | **Category** |
| --- | --- |
| Walks forward (number of steps) | Walk |
| Walks backward (number of steps) | Walk |
| Walks turning (number of steps) | Walk |
| Immobility (time while tortoise is immobile) | Immobility |
| Raises head and visual fixation (number of times) | Visual exploration |
| Turns head and visual fixation (number of times) | Visual exploration |
| Visual fixation (number of times) | Visual exploration |
| Smells the bottom of the box (number of times) | Olfactive exploration |
| Smells the sidewalls of the box (number of times) | Olfactive exploration |

We used a red box with a red bottom in order to put Hermann tortoises in a very new context (there were used to be in green boxes or in nature with green and brown colours). It has been demonstrated that they can distinguish these colours (Pelliteri-Rosa, 2010).

### Novel environment test (neophobia test)

Studying the tortoises’ behaviors in the 1h experiment we concluded that 10 minutes of observation were enough to visualize interindividual differences in exploration behavior, by consequence we conducted 10 minutes duration neophobia tests.

The experimental apparatus was a box of dimensions 80cm x 100cm and was divided into 12 cells of 26,5cm x 25cm

At the beginning of the experiment the tortoise was put under a hiding place which consisted in a grey plastic tunnel of dimensions 25cm x 25cm that covers the whole body of the tortoise (Figure 1)

**Figure 1:**
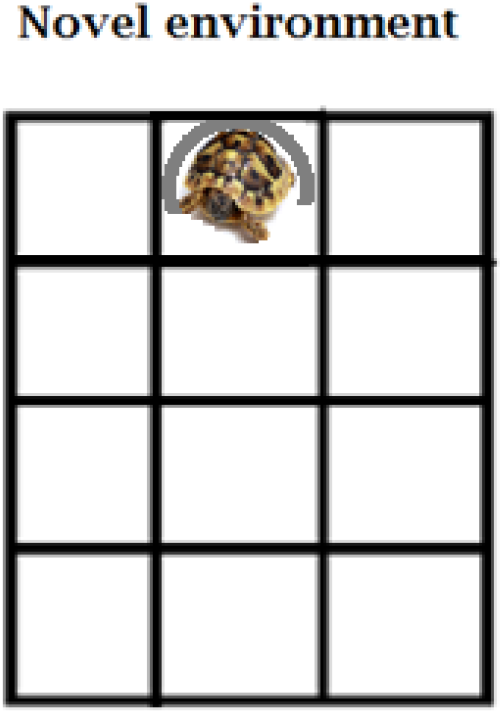
Aparatus, red wood box.

During the experiments we measured the time the tortoises took to exit the tunnel, the number of cells they visited (0 to 11) and we counted all the behaviors they expressed.

At the end of the experiment we cleaned the box in order to avoid potential effects of olfactive cues on the next individuals tested.

As we had measured the time of immobility and some tortoises went back into the tunnel we divided the number of times the tortoises walked by the remaining time in the box outside the tunnel and obtained the number of steps by minute:

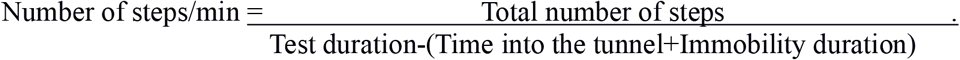

Three groups with different life experiences were tested:

- the Tortoise Village Group: 13 males and 13 females that have been raised by years at the Tortoise Village Center (they are tortoises that have been brought by people who cannot take care of them anymore or who found them on the road)
- the Chelonian Conservation Center Group: 12 males and 9 females that have been found in nature and brought to the center and who stay a short period in an isolated area not disturbed by public
- the Wild Field Group: 8 males and 13 females that live in wild conditions and that were tested in their natural field conditions.

For the tortoises that were raised in the Tortoise Village all the experiments were conducted putting the box into the enclosures. For the tortoises living into the wild the experimental box was brought to the field and the experiment was conducted in the place where the tortoise was found by telemetry.

We conducted Mann Whitney tests in each group in order to know if there were differences in behavior between males and females for each behavioral category.

We conducted Kruskal-Wallis tests and paired Mann Whitney tests between the three groups in order to know if they behave differently.

### Personalities

Individuals were grouped into two categories. Individuals that visited less than half of the cells (0 to 5 cells) were classified as shy (low explorers) and individuals that visited more than half of the apparatus cells (6 to 11 cells) were classified as bold (high explorers).

The box test was repeated two times for the tortoises of the conservation center group and the wild field group and it was repeated four times for the tortoise village group in order to see if the tortoises’ reactions were stable during time.

Tortoises were tested between 28/08/2015 and 10/09/2015 the first year and between 17/05/2016 and 24/05/2016 the second year. We choose those periods because temperature is higher and general activity of the Hermann’s tortoises is higher in this period and they all have finished their hibernation period. We had to concentrate in periods of the year when climate is similar and in sunny days while there was no rain and wind in order to avoid big differences in climate conditions between the tests.

### Antipredator reaction

Before the experiment in the box, the observer took the tortoise from its enclosure or from the field and noted its antipredator reaction. Tortoises expressed three kinds of reactions in this situation: retraction into the shell, nothing (the tortoise stays relaxed with the head and the legs outside the shell) or struggles (the tortoise moves her head and legs and tries to separate from the experimenter)

Chi square tests were conducted in order to see if there was a relation between the bold/shy individuals’ reaction in the neophobia test and the antipredatory test.

### Effect of personalities on wild exploration and dispersion

In previous studies of the team, the habitat use and home range of the wild Hermann Tortoises was analyzed (Sibeaux et al. 2016, Pille et al. 2017). Telemetric monitoring allowed us to measure the distances travelled by tortoises in the field during the days of monitoring over two periods in 2013 and 2014.

Tortoises were fitted with an AVM-K16 transmitter glued onto the shell. The transmitter plus resin represented <10% of individual body mass, a value assumed to be well tolerated in free-ranging tortoises (Lagarde et al., 2008). Each individual was located once a day, alternatively in the morning, around midday and in the afternoon. Coordinates were recorded using a Garmin GPS. Daily displacements of radio-tracked tortoises were calculated as the mean distance travelled per day.

19 tortoises (12 females and 7 males) of the wild field group described before were monitored by this way. We used the GPS coordinates obtained each day to compute the distances travelled by the tortoises.

We conducted Mann-Withney tests in order to know if there was a relation between the behavioral profile of the individual in the neophobia test and the day distances they travelled.

## Results

### Differences of behavior in the neophobia test

The comparison between the sexes in each group tested showed no differences for the different behaviors studied (Mann Whitney test p>0.05 for all the comparisons between sexes). By consequence, we could regroup females and males of each group for the following tests.

When we compared the three groups, we obtained differences in the expression of several behaviors (Figure 2). Tortoises from the Chelonian Research Center and from the Wild Field took significantly more time to exit the tunnel than the individuals from the Tortoise Village. Those tortoises explored significantly less squares in the experimental dispositive and stayed significantly more time in immobility than the tortoises from the Village.

**Figure 2.**
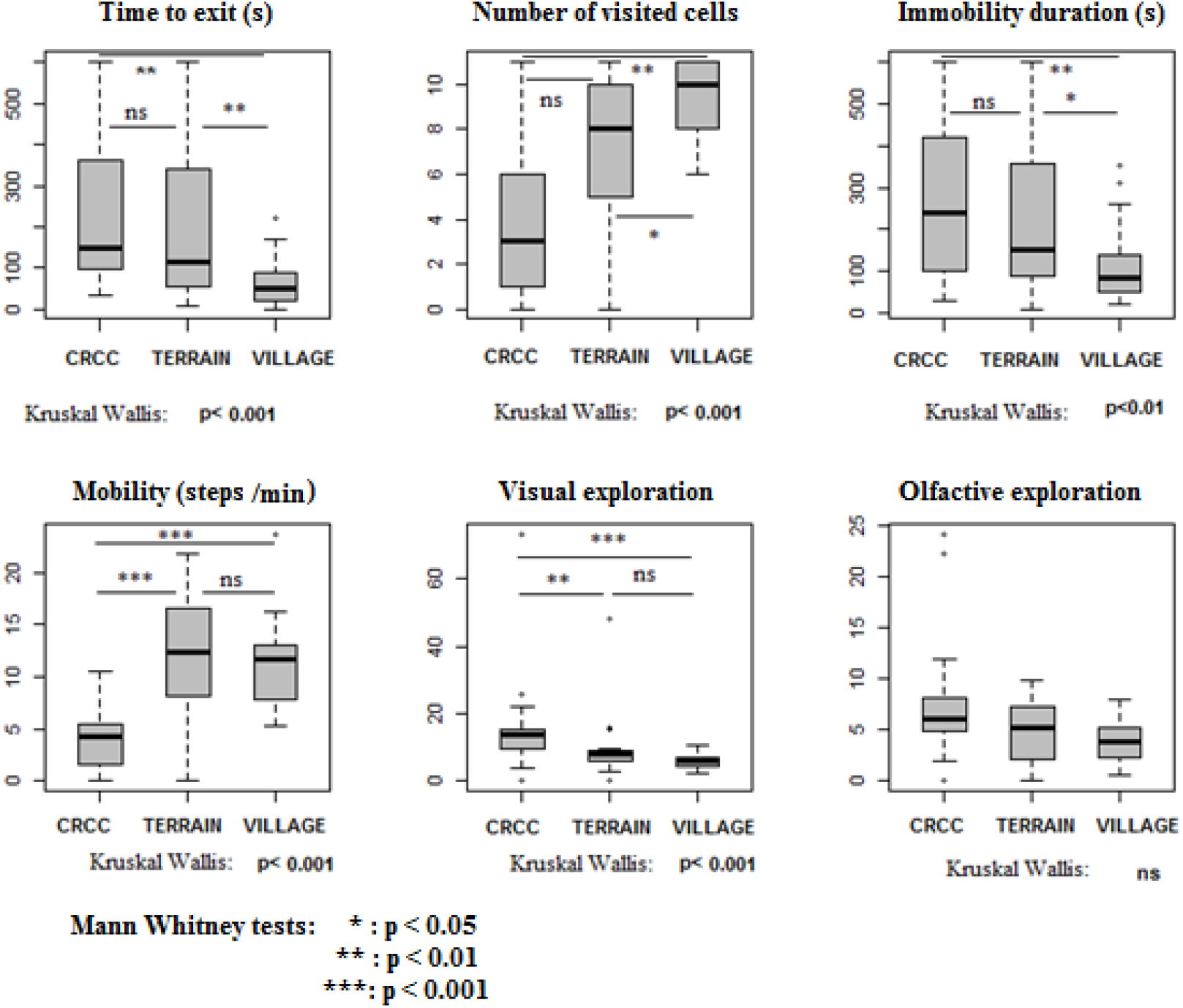
CRCC=Chelonian Research Center Terrain=Wild field Village=Tortoise Village.

During the exploration period, tortoises from the Field and from the Village elicit significantly more deambulation movements than the tortoises from the Chelonian Research Center. Tortoises from the Chelonian Research Center realized significantly more visual exploration of the dispositive than the tortoises from the Wild Field and from the Village. They were no differences in olfactive exploration between the three groups.

### Interindividual differences and personality

When we took all the data (the three groups in neophobia test and tests repetitions) and classified individuals into the bold (6 to 11 visited cells) and the shy (0 to 5 visited cells) categories we obtained different behavioral profiles (Figure 3). Bold individuals show significantly less time than the shy ones to exit the tunnel and they were significantly more mobile than the shy ones. Shy individuals tended to explore more the experimental dispositive in a visual way than bold ones (p=0,06) while bold individuals explored significantly more by olfactive exploration than shy ones (p=0.017).

**Figure 3:**
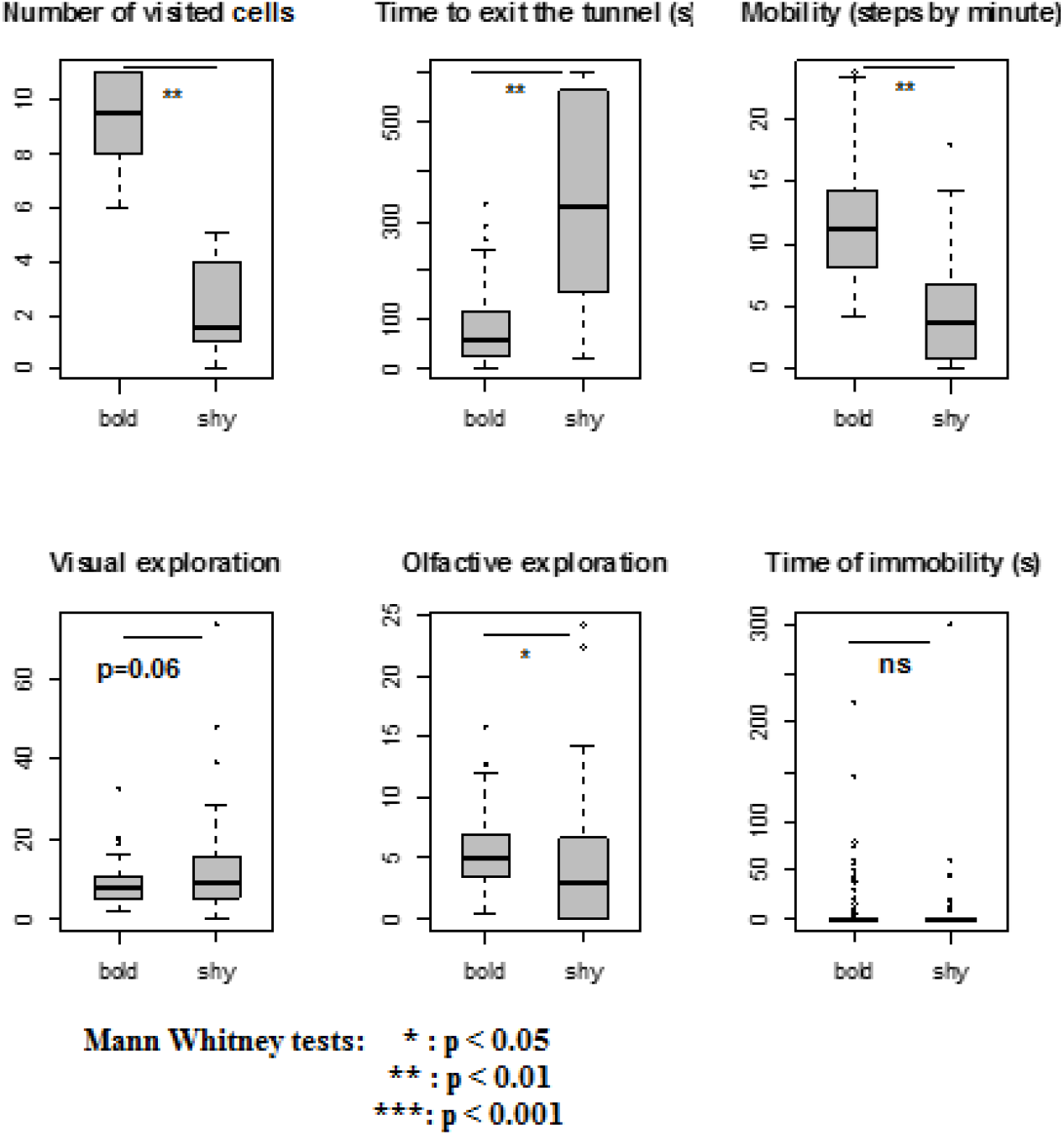
Behavioral characteristics of shy and bold individuals.

### Repeatability of explorative behavior

73,1% of the individuals behaved in the same way in the neophobia test and the repetition test. 26,9% of the individuals behave one time shy and one time bold. Chi square test conducted to a p value inferior to 0.01 indicating that individuals behave significantly more frequently in the same way (Figure 4a). However, half of the individuals that presented both behavioral profiles behave first in a shy way and then in a bold way, by consequence we can think that neophobia decreased in some individuals and conducted to a habituation of the apparatus.

**Figure 4:**
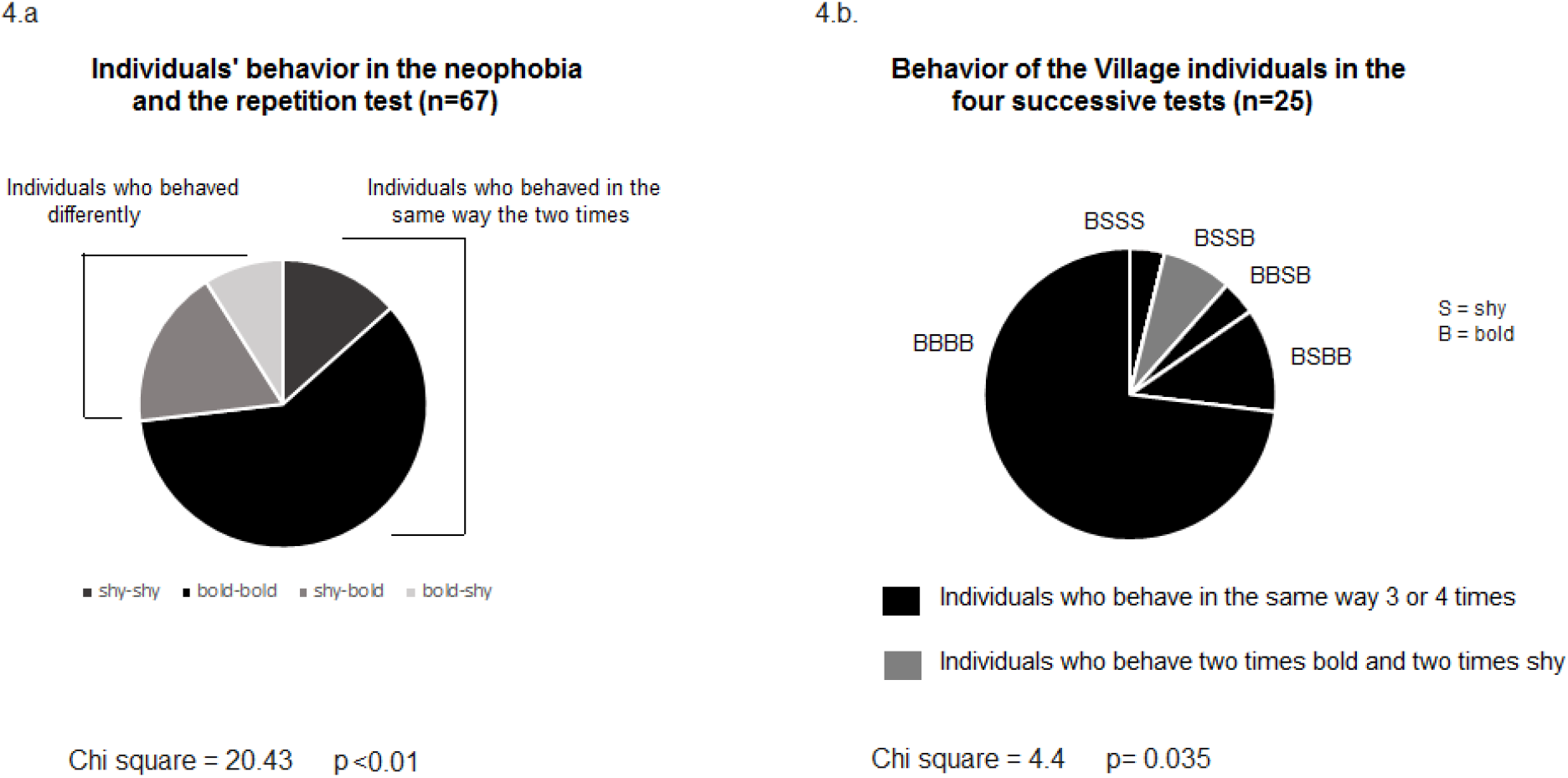
Behavior stability in the neophobia and repetition tests.

In the case of the individuals from the Tortoises’ Village who were tested four times, 92% of the individuals behave 3 or 4 times in the same way and 8% of the individuals behave twice bold and twice shy (Figure 4b). Chi square value (Chi=4.4, p=0.035) indicated that the observed proportions were significantly different from the proportions expected by hazard. This result also argues for a stability in the behavioral profiles.

### Antipredator reaction

We compared antipredatory reactions between males and females and obtained no differences (Figure 5a, Chi square=5.41, p=0.067).

**Figure 5:**
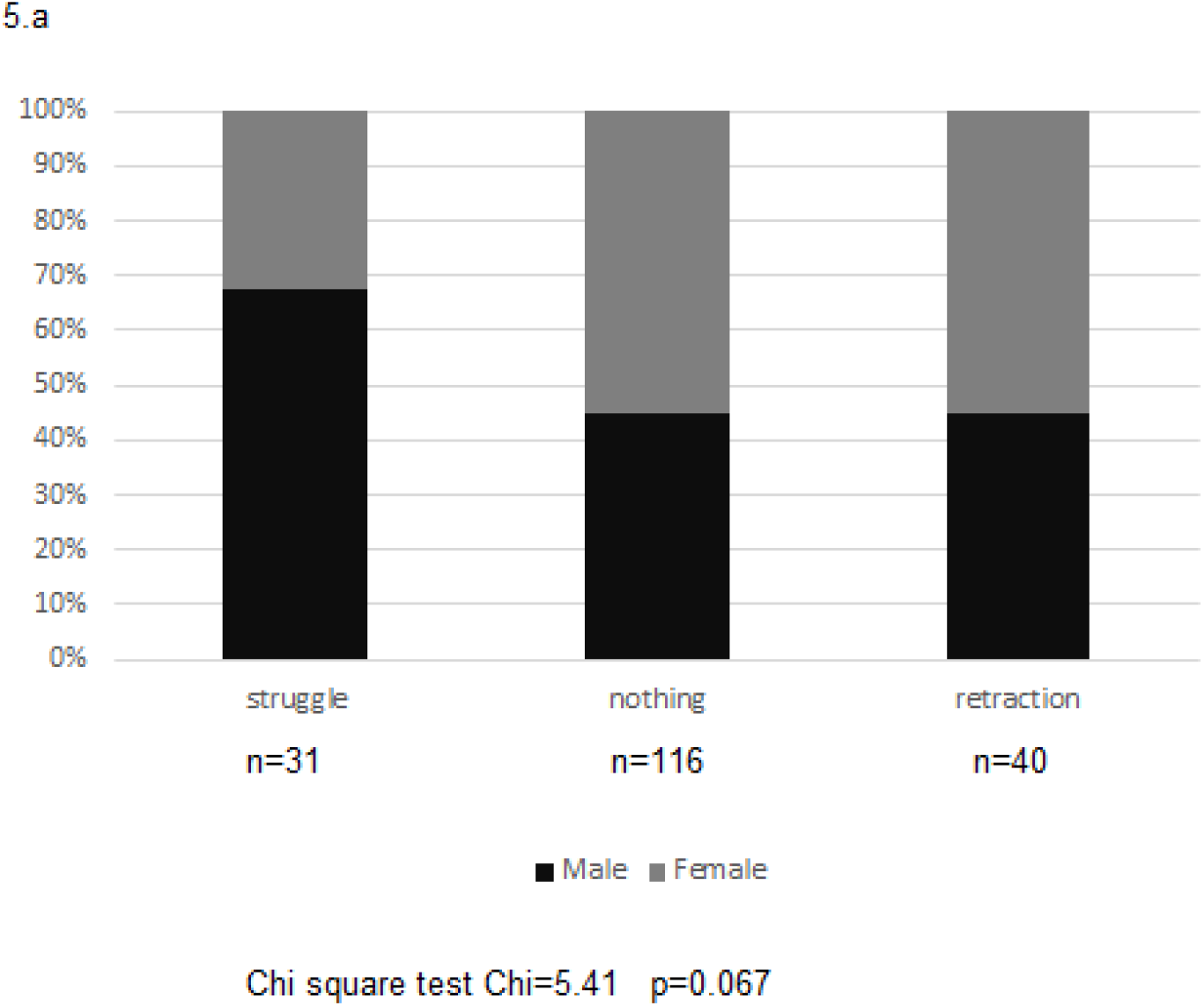
a. Antipredator behaviors and sex.

**b.**
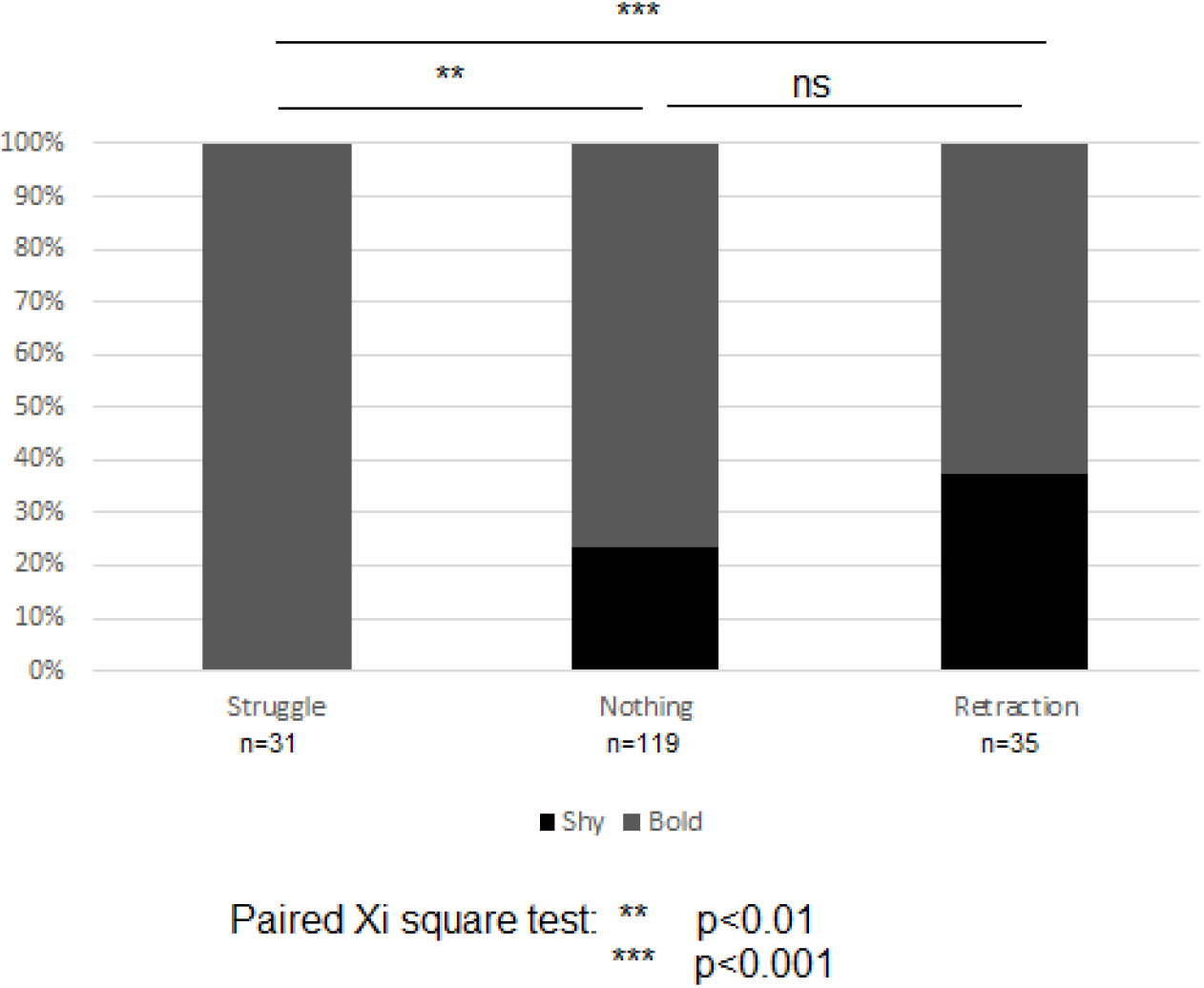
Antipredatory behaviors and behavioral profile (bold or shy) in the box.

**c.**
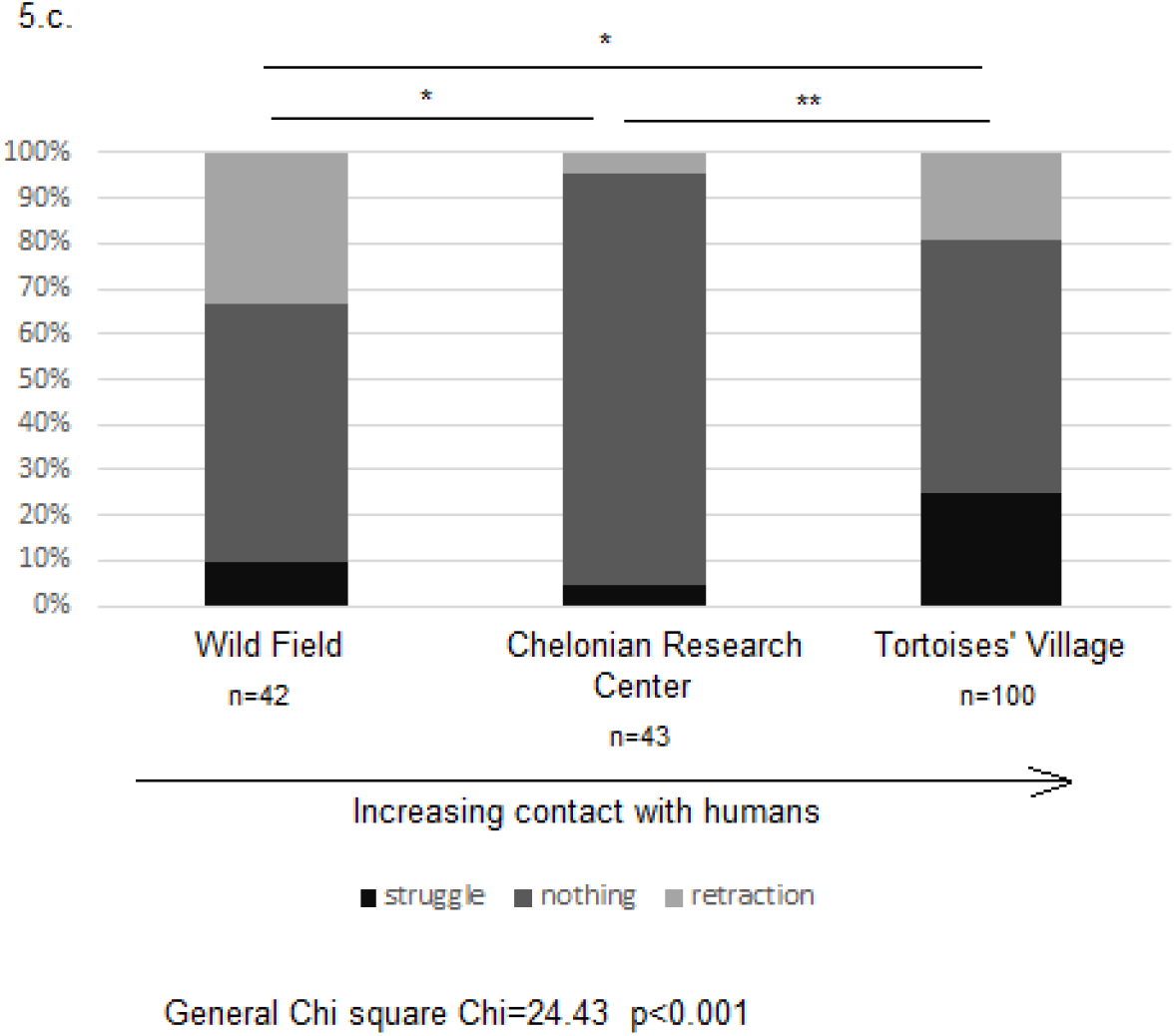
Antipredatory behaviors and frequency of contact with humans.

Concerning the comparison between different boldness types, when the tortoises were taken by the observer, only the individuals that behave in a bold way in the box struggled in the hand of the observer (Figure 5b). There were significantly more shy individuals in the relaxed group that did nothing and in the group that retracted into the shell than in the group that struggled. These observations indicates that nothing reaction and retraction are behaviors common to shy and bold individuals whereas struggle behavior is specific to bold individuals.

There were significant differences in antipredatory behavior between the groups with different histories and contact with humans. Individuals from the Tortoise Village struggled significantly more often than individuals from the Wild Field and from the Chelonian Research Center. Wild Field tortoises retracted significantly more than tortoises that are used to be manipulated by humans. Tortoises that have intermediate levels of manipulation (Chelonian Research Center group) presented significantly more often the intermediate relaxed antipredator reaction i.e. nothing.

### Neophobia and dispersion

Individuals that were shy in the neophobia test travelled daily distances significantly shorter than bold individuals (Figure 6, p <0.01). However there were no differences in the areas of the home ranges between these two groups (Figure 7b, p=0.73)

**Figure 7:**
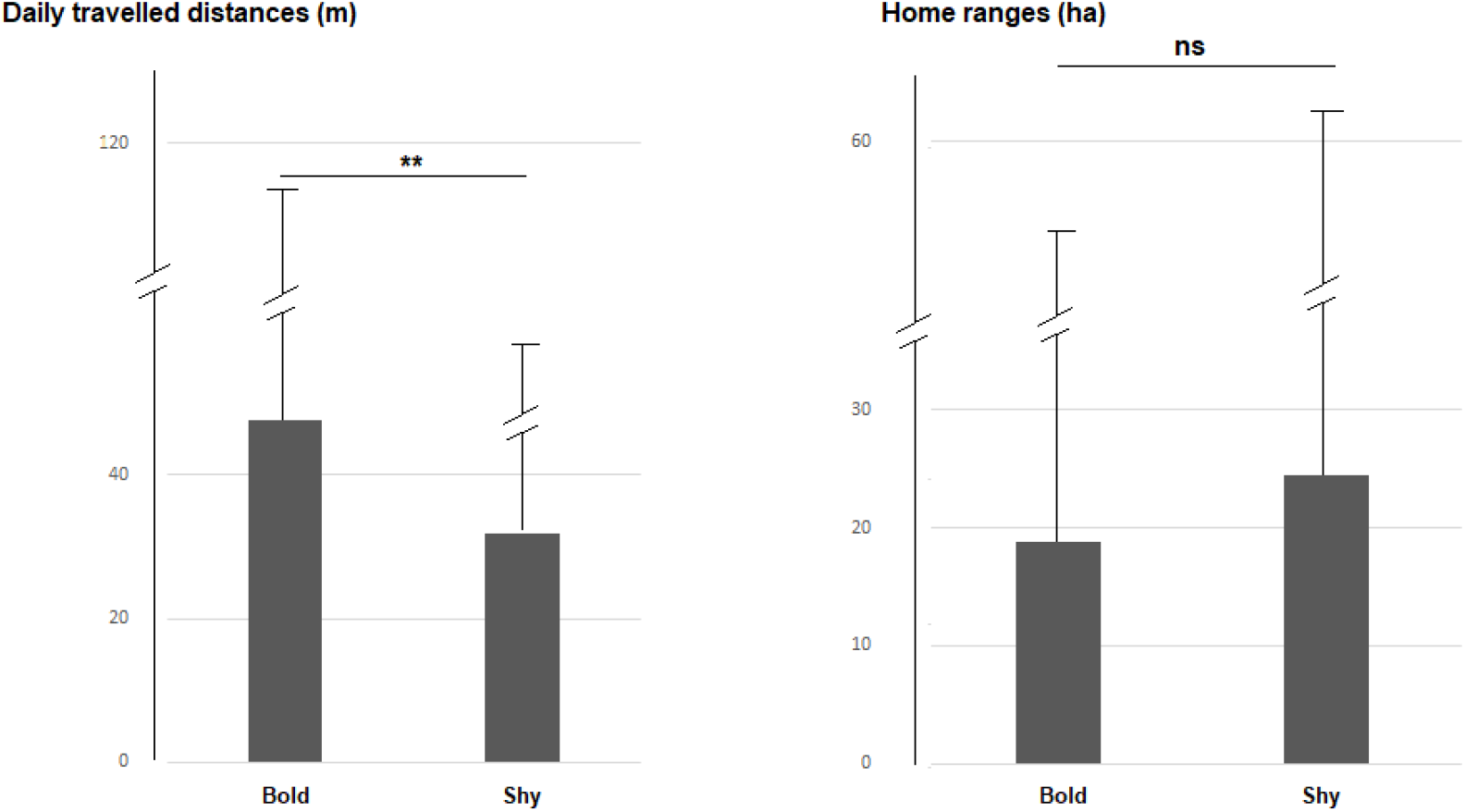
a. Daily distances traveled in 2013 and 2014 by bold and shy individuals. b. Homerange areas of bold and shy individuals during the years 2013 and 2014

